# Chronic Intermittent Ethanol Exposure Produces Sex- and Tissue-Specific Metabolomic Signatures Across the Gut-Liver Axis in Adult Mice

**DOI:** 10.64898/2026.06.14.732186

**Authors:** Julie Pollak, Reginald Cannady, Bo Wang, Antoniette M. Maldonado-Devincci

**Author notes:** To whom correspondence should be addressed: Antoniette M. Maldonado-Devincci, Ph.D., North Carolina A&T State University, Department of Psychology, 1601 East Market Street, New Science Building Room 360, Greensboro, North Carolina 27411, Bo Wang, Ph.D., Florida Institute of Technology, Department of Chemistry and Chemical Engineering, 150 West University Boulevard, Melbourne, FL 32901-6975. Funded by the 1R16GM149485 (AMD) and National Science Foundation 2306740 (BW).

## Abstract

Alcohol misuse leads to a range of health complications and induces various metabolic perturbations that impacts multiple physiological systems, including the cardiovascular system, liver, and gut microbiota. However, limited research has been reported on these metabolic profile changes, particularly using models of alcohol dependence such as after chronic intermittent ethanol (CIE) vapor exposure. This study investigated CIE-induced metabolomic alterations of CIE were investigated using fecal, liver, and serum samples of adult male and female C57BL/6J mice following 72 hr withdrawal. Significant metabolite changes were observed in both fecal and liver extracts and these changes were sex-specific. Both liver and fecal metabolites had systematic changes, while blood serum influences were limited after CIE. Female fecal samples showed higher metabolite perturbations than male samples according to PCA studies. The female samples showed significant butyrate downregulation and acetate upregulation, which are critical microbial products as beneficial microbe cell energy sources and influence intestinal absorption in the host. In addition, the female fecal samples showed significant downregulation of branched-chain amino acids including leucine, isoleucine, and valine, while male samples showed downregulation of glucose and taurine, with upregulated phenylalanine and tyrosine. In contrast, in the liver study, phenylalanine and tyrosine were upregulated while taurine was downregulated in females. Both sexes showed downregulation of liver glycine and glucose. These data indicate that CIE induces sex-specific metabolic perturbations in the mouse liver and fecal metabolome, and have implications for guy disturbances and liver damage observed following alcohol dependence. This study provides potential targets for future examination of mechanisms and treatment approaches for alcohol dependence.

## 1 Introduction

Alcohol consumption remains a widespread public health concern in the United States, affecting adolescents, adults, and older individuals and contributing substantially to preventable alcohol-related morbidity and mortality (CDC, 2025; Alcohol-Related Emergencies and Deaths in the United States, 2026a) [1]. Excessive alcohol use accounts for ∼178,000 deaths annually, with the majority attributable to chronic alcohol-related diseases rather than acute intoxication (CDC, 2025). Sustained alcohol exposure is causally linked to a number of pathologies, including alcohol-associated liver disease, pancreatitis, cardiovascular dysfunction, as well as multiple types of cancers including liver, breast, and gastrointestinal cancers (White et al., 2022; Alcohol-Related Emergencies and Deaths in the United States, 2026b; CDC, 2026). Alcohol-use disorders (AUD) remain highly prevalent in theUnited States (Grant et al., 2015). Importantly, alcohol use and dependence exhibit marked sex differences in adults, with males and females differing in drinking patterns, trajectories to dependence, and vulnerability to alcohol-related health consequences (Erol and Karpyak, 2015; Becker et al., 2017). Females in particular, often experience greater physiological and metabolic harm at lower levels of alcohol exposure compared to males and show a more rapid progression from initial use to dependence, increasing the risk for long-term adverse outcomes associated with sustained high alcohol use (Erol and Karpyak, 2015; Becker et al., 2017).

Sex differences in AUD trajectories and alcohol-related disease burden are well documented in the clinical literature. Although, historically AUD levels have been higher among men, similar rates of elevated use and associated adverse health outcomes have rapidly grown in women over the last decade, resulting in a narrow gap between sexes (Karaye et al., 2023). Importantly, women experience these adverse health outcomes at lower levels of exposure, including increased vulnerability to alcohol-associated liver disease, cardiovascular dysfunction, and metabolic impairment (Baraona et al., 2001; Piano et al., 2020; Singal and Mathurin, 2021; Greaves et al., 2022; Llamosas-Falcón et al., 2022). These sex-dependence health risks are partly attributable to differences in pharmacokinetics and systemic alcohol exposure, including reduced first passed metabolism, higher blood ethanol concentrations, differences in body composition, and ovarian hormone-dependent modulation of ethanol clearance and tissue responses (Baraona et al., 2001; Greaves et al., 2022). Together, these indicate that sex is a critically important factor implicated in alcohol-related disease risk and progression.

Chronic intermittent ethanol (CIE) exposure is a well-established model used to mimic repeated cycles of ethanol intoxication and withdrawal that produce dependence-like phenotypes relevant to AUD (Becker and Lopez, 2004; Becker, 2014) (Ferguson et al., 2022). This exposure paradigm has been shown to induce robust and persistent molecular adaptations across central and peripheral tissues, including alterations in gene expression and physiological systems that extend beyond the brain (Lundqvist et al., 1994; Becker, 2017; Liu and Crews, 2017; Macht et al., 2020; Ferguson et al., 2022; Shobande et al., 2026). Despite extensive use of the CIE model, to characterize neurobiological and transcriptional consequences of chronic alcohol exposure, the mechanisms through which CIE drives systemic metabolic disruption, particularly in a sex-specific and organ-specific manner, remain incompletely understood (Shobande et al., 2026).

Preclinical studies that include both males and females consistently demonstrate that CIE and similar intermittent or binge-pattern exposure paradigms result in sex-specific effects on alcohol intake, withdrawal-associated behaviors, and peripheral metabolic profiles (Jury et al., 2017; Rivera-Irizarry et al., 2023; Shobande et al., 2026). In C57BL/6J mice, CIE robustly increases voluntary ethanol drinking in males, whereas females consistently fail to show escalations in CIE-induced drinking, despite comparable and often higher baseline ethanol drinking levels (Jury et al., 2017). Beyond alcohol consumption, males and females differ in sensitivity to ethanol-induced sedation, withdrawal-associated nociception, and affective behaviors during both early and protracted withdrawal, with sex-specific differences in both magnitude and temporal profile across paradigms (Jury et al., 2017; Brandner et al., 2023; Rivera-Irizarry et al., 2023). Extending beyond behavioral endpoints, intermittent ethanol exposure paradigms during adolescence produces sex-specific alterations in peripheral metabolic profiles, with males demonstrating more pronounced short-term disruptions and females showing distinct time-dependent patterns across withdrawal intervals (Shobande et al., 2026). Collectively, these findings provide important context for examining whether similar sex-specific metabolic adaptations emerge following CIE exposure in adulthood.

Ethanol exposure engages the gut-liver axis as an important contributor to peripheral metabolic changes during withdrawal (Szabo, 2015; Bajaj, 2019; Albillos et al., 2020). Ethanol consumption is consistently associated with changes in the gut environment that alter the profile of gut-associated metabolites, including short-chain fatty acids, amino acid-related metabolites, and bile acids, which are reflected in peripheral metabolic profiles following ethanol exposure (Bajaj, 2019; Collins et al., 2023; Hsu and Schnabl, 2023). In preclinical CIE models, withdrawal is associated with peripheral metabolic alterations at the level of gut-associated metabolites, hepatic metabolic processing, and circulation profiles, supporting investigation of fecal, liver, and serum profiles within the same withdrawal window (Bajaj, 2019; Albillos et al., 2020; Tilg et al., 2022; Hsu and Schnabl, 2023; Shobande et al., 2026).

The advancements in metabolomic studies provide promising potential in investigating the underlying molecular level changes after CIE (Hwang et al.). Metabolites are the chemical intermediates or final products of the metabolism and provide a functional readout of ongoing biological processings, integrating genetic, physiological, and environmental influences, including ethanol exposure (Mamas et al., 2011; Ivanisevic and Thomas, 2018; Bajaj, 2019; Tilg et al., 2022; Hsu and Schnabl, 2023). By examining combinational variation of metabolites, metabolomic approaches capture coordinated pathway-level changes that reflect alcohol– and withdrawal-related perturbations not apparent from individual metabolites alone (Bajaj, 2019; Tilg et al., 2022). Severe thiamine deficiency has been identified in serum samples from individuals with alcoholic brain disease, with clinically meaningful (Martin et al., 2003) symptom improvement observed following thiamine administration in mixed clinical populations (Mulholland et al., 2005). These patient samples demonstrated the utility of peripheral metabolite measures for assessing treatment efficacy and tracking metabolic changes following intervention (Tallaksen et al., 1992). Metabolomics studies have also been used to evaluate pharmacological interventions aimed at reversing alcohol-induced metabolic dysregulation across peripheral tissues (Harrigan et al., 2008; Bajaj, 2019; Tilg et al., 2022). Metabolic profiling following alcohol intake and alcohol dependence have revealed disruptions in central energy metabolism and amino acid-related pathways (Harada et al., 2016; Irwin et al., 2018; Zhu et al., 2021) and nuclear magnetic resonance (NMR)-based approaches has identified alcohol-related alterations in redox balance and metabolites linked to energy metabolism (Irwin et al., 2018).. In addition, alcoholic liver disease and liver injury have been investigated using metabolomics approaches, highlighting the value of liver-based metabolic profiling for assessing alcohol-related metabolic disruption at a systems level (Liang et al., 2015; Ganesan and Suk, 2022; Jaber et al., 2023).

Despite these advantages, relatively few studies have examined metabolic alterations following CIE exposure, with limited attention to sex-by-tissue differences and limited integration across gut, liver, and serum. This gap likely reflects the challenge of capturing dynamic, systems-level metabolic processes rather than single steady states (Deneke and Di Talia). Metabolomic analyses provide an approach for examining metabolic alterations across gut, liver, and circulating levels following CIE exposure.

This study used NMR-based metabolomics to characterize sex– and tissue-specific metabolic alterations in fecal, liver, and serum samples from adult C57BL/6J mice following CIE exposure. We aimed to identify candidate metabolites and metabolic pathways associated with CIE and to determine whether metabolic signatures differ by sex and tissue. We hypothesized that CIE would produce sex– and tissue-specific metabolomic alterations and that fecal metabolic profiles may reflect gut-associated microbiome-linked changes.

## 2 Methodology

### 2.1 Animals study

Adult male and female C57BL/6J mice (about 2-3 months old; n=10-13 per group) were derived from in-house established breeding pairs and weaned on postnatal day (PND 21) with same-sex littermates. Mice were maintained in the animal colony under standard housing conditions, including corn cob bedding, nestlet, group housing, routine cage maintenance, and daily health monitoring, until inclusion in the study. Once animals were assigned to experimental conditions, animals were handled daily to allow acclimation to experimenter manipulation. Mice were group-housed (4-5 per cage) with *ad libitum* access to food and water throughout the air or ethanol exposure periods. All mice were housed in a temperature– and humidity-controlled room on 12-hr light/dark cycles with lights on from 0700-1900 hr. Body weights were recorded at each chamber exposure. All procedures were conducted in accordance with the National Institutes of Health Guidelines and were approved by the North Carolina Agricultural and Technical State University Institutional Animal Care and Use Committee.

### 2.2 Ethanol exposure experiment

Mice were exposed to repeated intermittent air (AIR control group) or ethanol vapor (CIE group) across three exposure cycles. Each cycle consisted of four consecutive days of exposure followed by three days off, during which mice remained in their homecage except for regular cage maintenance.

On Day 1 of each exposure cycle, mice were weighed and administered an intraperitoneal injection (0.02 mL/g) of pyrazole (1 mmol/kg), an alcohol dehydrogenase inhibitor, used to stabilize blood ethanol concentrations. AIR control mice received pyrazole combined with saline, whereas CIE mice received pyrazole combined with ethanol (1.6 g/kg; 8% w/v). Immediately following injection, mice were placed into the inhalation chambers (23 in x 23 in x 13 in; Plas Labs, Lansing, MI) at ∼1700 hr and remained in the chambers for 16 hours overnight (∼1700 to 0900 hr).

Inhalation chambers were supplied with either room air (AIR group) or volatilized ethanol vapor (CIE group) delivered at a rate of 10 L/min. Ethanol vapor was generated by passing air through an air stone (gas diffuser) submerged in 95% ethanol. These exposure parameters match previously validated CIE vapor conditions in C57BL/6J mice that reliably produce blood ethanol concentrations within the target range of approximately 150-275 mg/dL, as demonstrated in prior studies using identical or highly comparable exposure conditions (Griffin et al., 2009; Lopez et al., 2012; Maldonado-Devincci and Cook, 2014; Zamudio et al., 2021) throughout the exposure period.

Blood ethanol concentrations were not directly assessed in the present cohort to minimize repeated blood sampling-associated stress and to preserve circulating metabolites for downstream serum-based analyses, as both handling and blood collection procedures can acutely alter circulating metabolite profiles in mice (Lee et al., 2023; Papageorgiou et al., 2024). This consideration is particularly relevant when serum metabolomics is a primary experimental outcome, as many factors related to sample collection, handling, and physiological stressors can contribute to variability in blood-based metabolomic measurements (Emwas et al., 2025). Instead, exposure validity was supported through the use of previously validated CIE exposure parameters established in our prior work and in the broader CIE literature using C57BL/6J mice, where blood ethanol concentrations were measured under comparable exposure conditions (Maldonado-Devincci et al., 2014, 2016; Zamudio et al., 2021).

Mice were removed from the chambers at approximately0900 hr and returned to their home cages. Animals remained undisturbed for three days between exposure cycles. Mouse fecal, serum, and liver samples were collected three days after completion of the final AIR or CIE exposure cycle.

### 2.3 Metabolite extraction

Three different sample types were analyzed including fecal, liver tissue, and serum. Separate extraction procedures were used for each sample type to account for differences in physical and biochemical properties (Moosmang et al.). For all sample types, approximately 100 mg of material was used for consistency and accuracy.

For fecal samples, a water extraction was carried out from a dry and frozen state using previously established methods (Gratton et al., 2016). Samples were centrifuged at 4°C at 14,800 rpm for 15 minutes. The supernatant was transferred to a clean tube. Supernatants were then mixed with a phosphate buffer of deuterium oxide (D_2_O) to a final composition of 10% D_2_O, 0.1 M phosphate buffer (pH 7.4), and 0.5 mM trimethylsilyl propanoic acid (TSP). The final solution was centrifuged again at the same conditions and transferred to 5 mm NMR tubes.

Liver tissue samples were extracted using a two-step polar extraction method as previously described (Wu et al., 2008). Briefly, tissue homogenized in ice-cold methanol-water (2.5:1 v/v), followed by extraction with ice-cold chloroform:water (1:1, v/v). The upper polar phase was collected and dried using a Savant SpeedVac Vacuum System. Dried extracts were resuspended in the same D_2_O phosphate buffer described above, with the same final concentrations of TSP, D_2_O, and phosphate buffer.

Serum samples were extracted using an NaCl-based method adapted from a previously reported protocol (Glader et al., 2019). Samples were treated with a 0.9% NaCl solution and subsequently mixed with the same D_2_O phosphate buffer described above, yielding identical final concentrations of TSP, D_2_O, and phosphate buffer.

### 2.4 NMR metabolomics analysis

Nuclear magnetic resonance (NMR) analysis was performed using a Bruker Ascend 400 MHz high-resolution spectrometer equipped with an Xpress autosampler. Data acquisition was conducted with ICON-NMR software (Bruker Biospin). Spectra were phased and calibrated to TSP using Bruker Topspin 4.06 (Bruker Biospin).

Metabolomic bucketing methods were carried out using AMIX (Bruker Biospin) with a bucketing approach previously reported (Wang et al., 2020). Metabolite identification was conducted using Chenomx 8.6 (Chenomx Inc). Principal component analysis (PCA) and biplots were generated using Pareto scaling and mean centering in PLS Toolbox (Eigenvector, Inc) based on MATLAB (MathWorks, Inc). Pathway analysis was performed using the KEGG database and MetaboAnalyst version 4.0.

### 2.5 Design and Analyses

Fold changes were calculated as the mean metabolite concentrations in CIE samples divided by the mean metabolite concentration in the AIR group. Fold change values greater than 1 indicated metabolite upregulation, whereas fold change values less than 1 indicated metabolite downregulation. Group comparisons were assessed using two-way ANOVA with sex (2; Male, Female) and Exposure (2; CIE, AIR) as between subject factors and planned within-sex t-test comparisons, with statistical significance denoted by a p-value of less than 0.05.

## 3 Results

### 3.1 Metabolomic response of fecal extracts after CIE

In the fecal extracts, a total of 42 metabolites were obtained as shown in **Table 1**. The metabolite distributions in the PCA studies, shown in **Figure 1**, revealed partial separation between the mice subjected to ethanol-CIE and the control-AIR in both male and female groups. However, the PCA score plot for female extracts showed a better visual separation when compared to the male samples judging by the distance between the center of the control cluster in red and CIE group cluster in green as indicated in **Figure 1**. PCA describes the major combinational contributions of all metabolites in the study between different groups, therefore, the major metabolites perturbation was observed after ethanol treatment in both male and female mice. Moreover, the male and female samples showed different responses in the metabolite profiling. In the male samples, the metabolites that were significantly upregulated were aspartate, methionine, nicotinate, phenylalanine, tyrosine, tryptophan, and alanine (p<0.05). Whereas glucose, glycocholate, and taurine were significantly downregulated (p<0.05). On the other hand, key metabolites that were upregulated in the female fecal samples were acetate, asparagine, histidine, trimethylamine, tryptophan, methylamine, and nicotinate (p<0.05). Metabolites such as glycocholate, butyrate, and many amino acids such as leucine, isoleucine, valine, and alanine were significantly downregulated in the female fecal extracts (p<0.05). Only three metabolites, tryptophan, nicotinate, and glycocholate were significantly changed in the same direction in both males and females (out of 11 in males and 16 in females).

**Figure 1:**
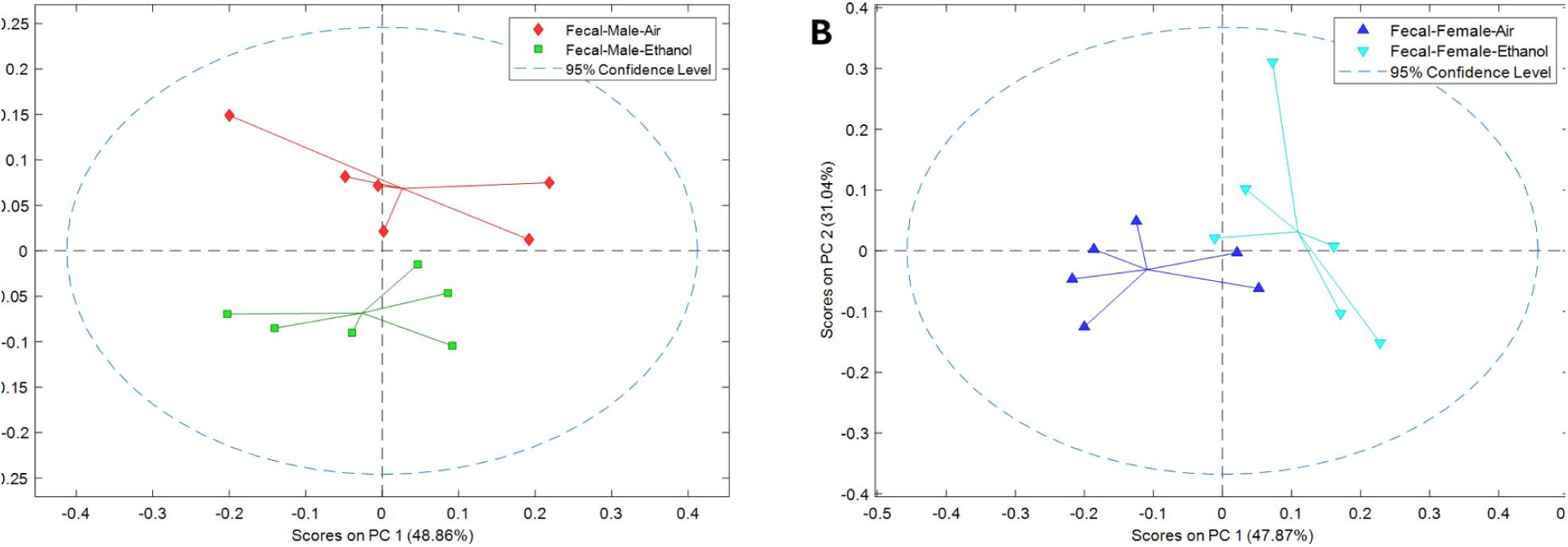
PCA studies on ethanol fecal metabolites vs. control. A is a score plot for the male samples and B is for the female samples. Pareto Scaling was applied.

**Table 1:** The fold change and p values of fecal metabolites obtained from male and female samples. Two-way ANOVA analyses were applied for Sex and Exposure and the significant interaction F and p-values are highlighted in bold (p<0.05).

|  | Male fecal | Female fecal | Two-way Interaction ANOVA |  |
| --- | --- | --- | --- | --- |
| Metabolites | Fold change | Fold change | F value | p-value |
| 1,3-Dihydroxyacetone | 1.09 | 1.05 | (1, 20) = 0.2195 | 0.6445 |
| 2-Oxoisocaproate | 1.00 | 1.17 | F (1, 20) = 3.651 | P=0.0705 |
| 3-Methyl-2-oxovalerate | 1.06 | 1.01 | F (1, 20) = 0.2066 | P=0.6544 |
| Acetate | 0.97 | 1.16 | F (1, 20) = 3.001 | P=0.0986 |
| **Alanine** | **1.09** | **0.83** | **F (1, 20) = 12.82** | **P=0.0019** |
| Arabinose | 0.92 | 0.97 | F (1, 20) = 0.1818 | P=0.6744 |
| **Asparagine** | **0.88** | **1.31** | **F (1, 20) = 10.54** | **P=0.0040** |
| **Aspartate** | **1.28** | **0.93** | **F (1, 20) = 12.54** | **P=0.0020** |
| Butyrate | 0.96 | 0.89 | F (1, 20) = 0.8093 | P=0.3790 |
| Choline | 1.04 | 1.07 | F (1, 20) = 0.1804 | P=0.6755 |
| Ethanol | 0.94 | 0.91 | F (1, 20) = 0.6179 | P=0.4411 |
| **Ethanolamine** | **1.09** | **0.84** | **F (1, 20) = 7.938** | **P=0.0106** |
| Formate | 0.86 | 8.65 | F (1, 20) = 3.357 | P=0.0819 |
| Fumarate | 1.43 | 1.63 | F (1, 20) = 0.1377 | P=0.7145 |
| Glucose | 0.79 | 0.99 | F (1, 20) = 2.882 | P=0.1051 |
| Glutamate | 1.04 | 0.98 | F (1, 20) = 1.317 | P=0.2647 |
| Glutamine | 1.06 | 0.95 | F (1, 20) = 1.231 | P=0.2803 |
| Glycine | 1.02 | 0.91 | F (1, 20) = 3.135 | P=0.0919 |
| Glycocholate | 0.53 | 0.46 | F (1, 20) = 0.9911 | P=0.3314 |
| **Histidine** | **0.88** | **1.44** | **F (1, 20) = 9.420** | **P=0.0061** |
| Hypoxanthine | 1.05 | 1.19 | F (1, 20) = 1.206 | P=0.2852 |
| **Isoleucine** | **1.06** | **0.74** | **F (1, 20) = 11.46** | **P=0.0029** |
| Lactate | 1.12 | 1.15 | F (1, 20) = 0.07122 | P=0.7923 |
| **Leucine** | **1.03** | **0.71** | **F (1, 20) = 20.10** | **P=0.0002** |
| Methanol | 0.99 | 1.02 | F (1, 20) = 0.3144 | P=0.5812 |
| Methionine | 1.16 | 1.00 | F (1, 20) = 3.205 | P=0.0886 |
| **Methylamine** | **0.99** | **1.24** | **F (1, 20) = 4.999** | **P=0.0369** |
| Nicotinate | 1.25 | 1.22 | F (1, 20) = 0.1182 | P=0.7346 |
| **Phenylalanine** | **1.31** | **0.91** | **F (1, 20) = 17.80** | **P=0.0004** |
| Propionate | 1.02 | 0.82 | F (1, 20) = 4.272 | P=0.0519 |
| **Putrescine** | **0.96** | **0.81** | **F (1, 20) = 8.221** | **P=0.0095** |
| Sarcosine | 1.05 | 1.00 | F (1, 20) = 0.5867 | P=0.4526 |
| Succinate | 0.98 | 1.12 | F (1, 20) = 1.462 | P=0.2407 |
| Taurine | 0.78 | 1.05 | F (1, 20) = 3.210 | P=0.0883 |
| Threonine | 1.04 | 0.98 | F (1, 20) = 0.6203 | P=0.4402 |
| **Trimethylamine** | **0.93** | **1.56** | **F (1, 20) = 10.17** | **P=0.0046** |
| Tryptophan | 1.46 | 1.20 | F (1, 20) = 3.080 | P=0.0945 |
| **Tyrosine** | **1.49** | **0.96** | **F (1, 20) = 16.04** | **P=0.0007** |
| Uracil | 2.13 | 7.52 | F (1, 20) = 0.6123 | P=0.4431 |
| **Valine** | **1.15** | **0.75** | **F (1, 20) = 13.44** | **P=0.0015** |
| Xanthine | 0.94 | 1.16 | F (1, 20) = 1.935 | P=0.1795 |
| Xylose | 0.90 | 0.98 | F (1, 20) = 0.3657 | P=0.5522 |

Individual metabolites depicted in Figure 2 showed statistically significant Sex by Exposure interactions (Table 1) or significant main effects of ethanol. Specifically, female AIR mice had higher metabolite levels than control males and ethanol reduced metabolite levels compared to female controls alanine (Figure 2A; p = 0.0002, p = 0.0019), valine (Figure 2C; p = 0.0007, p = 0.0018), leucine (Figure 2D; p < 0.0001, p < 0.0001), ethanolamine (Figure 2F; p = 0.0002, p = 0.01), isoleucine (Figure 2G; p = 0.001, p = 0.0008), and putrescine (Figure 2H; p = 0.0017, p < 0.0001). Ethanol exposure reduced metabolite levels in both males and females as supported by a main effect of Exposure for glycocholate [Figure2B, (F (1, 20) = 48.17), p < 0.0001] and nicotinate [Figure 2J, (F (1, 20) = 19.76), p = 0.0002]. CIE increased aspartate levels in males (p = 0.0009) and these levels were higher compared to similarly exposed CIE females (p = 0.01; Figure 2E). AIR male histidine levels were higher compared to AIR females (p = 0.0278) and that CIE increased histidine levels in females compared to AIR females (p = 0.0044; Figure 2I). CIE increased phenylalanine levels in males (p = 0.0003) and these levels were higher compared to similarly exposed CIE females (p = 0.0154; Figure 2K). Tryptophan levels were higher in CIE males compared to AIR males (p = 0.0006) and CIE females (p = 0.0045; Figure 2L). Whereas asparagine levels were higher in CIE females compared to AIR females (p = 0.0057) and CIE males (p = 0.0191; Figure 2M). Higher methylamine levels were observed in CIE females compared to AIR females (p = 0.0006) and no CIE effects in males (Figure 2N). Figure 2O shows a higher sex difference in trimethylamine with higher levels in AIR males compared to AIR females (p = 0.0029) and CIE increased levels in females compared to their AIR controls (p = 0.0011). Finally, CIE increased tyrosine levels in males compared to AIR male mice (p < 0.0001) and were higher than CIE females (Figure 2P; p = 0.0019).

**Figure 2:**
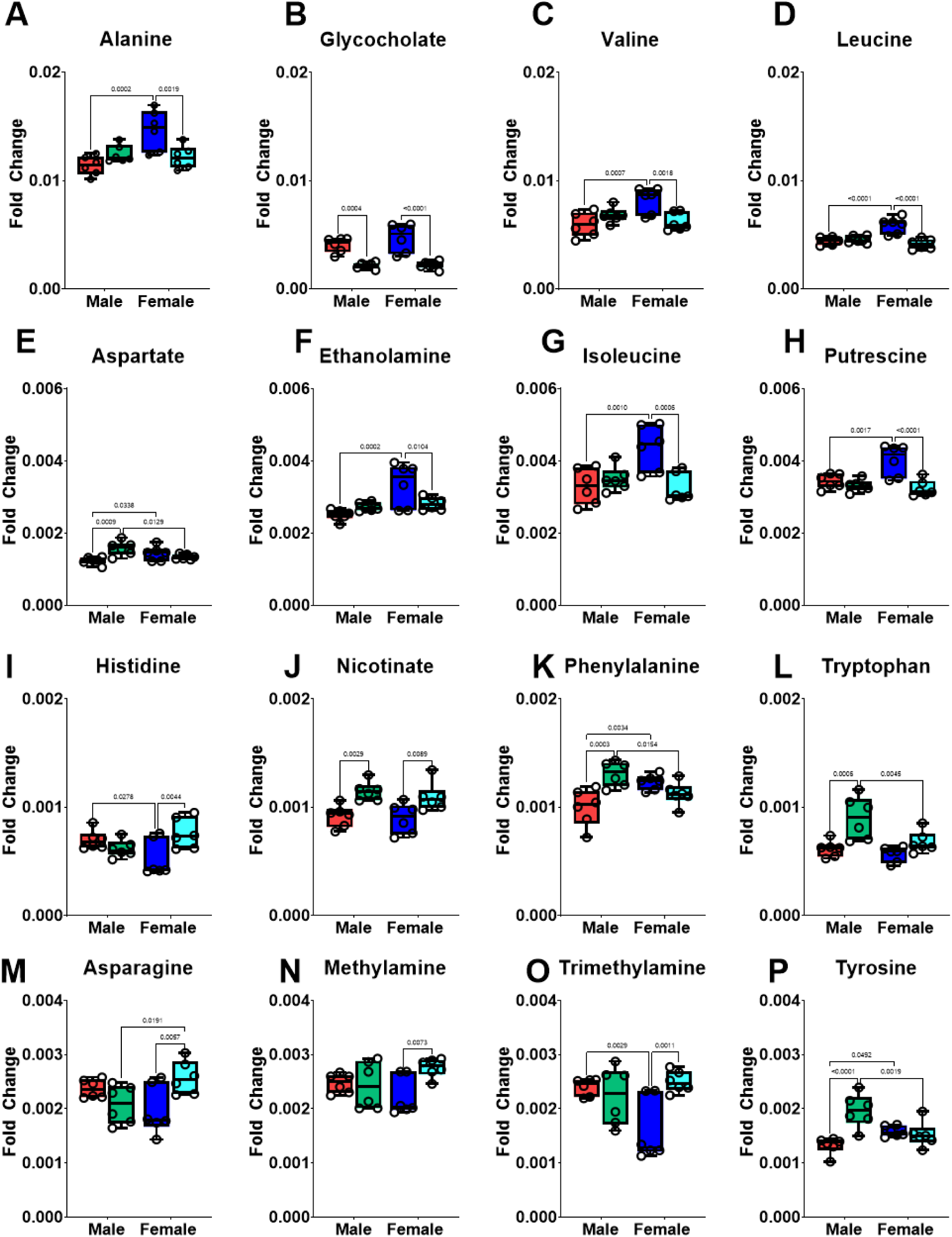
Fecal metabolite comparison of individual metabolites following Chronic Intermittent Ethanol (CIE) exposure. Individual panels (A-P) show the relative concentration of fecal metabolites that exhibited a significant Sex and Exposure interaction or a significant main effect of Exposure. Groups are color-coded as follows: Red bars represent AIR (control) males; Green bars represent CIE males; Blue bars represent AIR (control) females; and Teal bars represent CIE females. Specific p-values for all statistical comparisons are detailed in Section 3.1.

### 3.2 Metabolomic response of liver extracts after CIE

In the liver extracts, 44 metabolites were observed as shown in **Table *2***. The metabolite distributions in the PCA studies, shown in Figure 3, revealed a distinct difference between the groups subjected to CIE vs. AIR in both male and female groups. The female PCA score plot showed a clear partial separation between the AIE and CIE samples when compared to that of the male samples when visually comparing the cluster centers. Key metabolites that were significantly upregulated in the male liver samples were histidine, o-phosphocholine, fumarate, glutathione, β-Alanine, inosine, and uracil. Whereas glycine and isoleucine were downregulated in the male liver samples (p<0.05). In contrast, 17 metabolites were significantly changed in the female samples. The key metabolites that were upregulated were citric acid, formate, histidine, methionine, tyramine, tyrosine, and uracil (p<0.05). Metabolites such as choline, glycine, maltose, o-phosphocholine, and taurine were all downregulated in the female liver extracts (p<0.05). Female samples had 17 noteworthy metabolite changes, while male samples had 10 metabolite changes that were statistically significant, with 7 shared metabolites (Formate, Fumarate, Glucose, Glycine, Histidine, O-Phosphocholine and Uracil).

**Figure 3:**
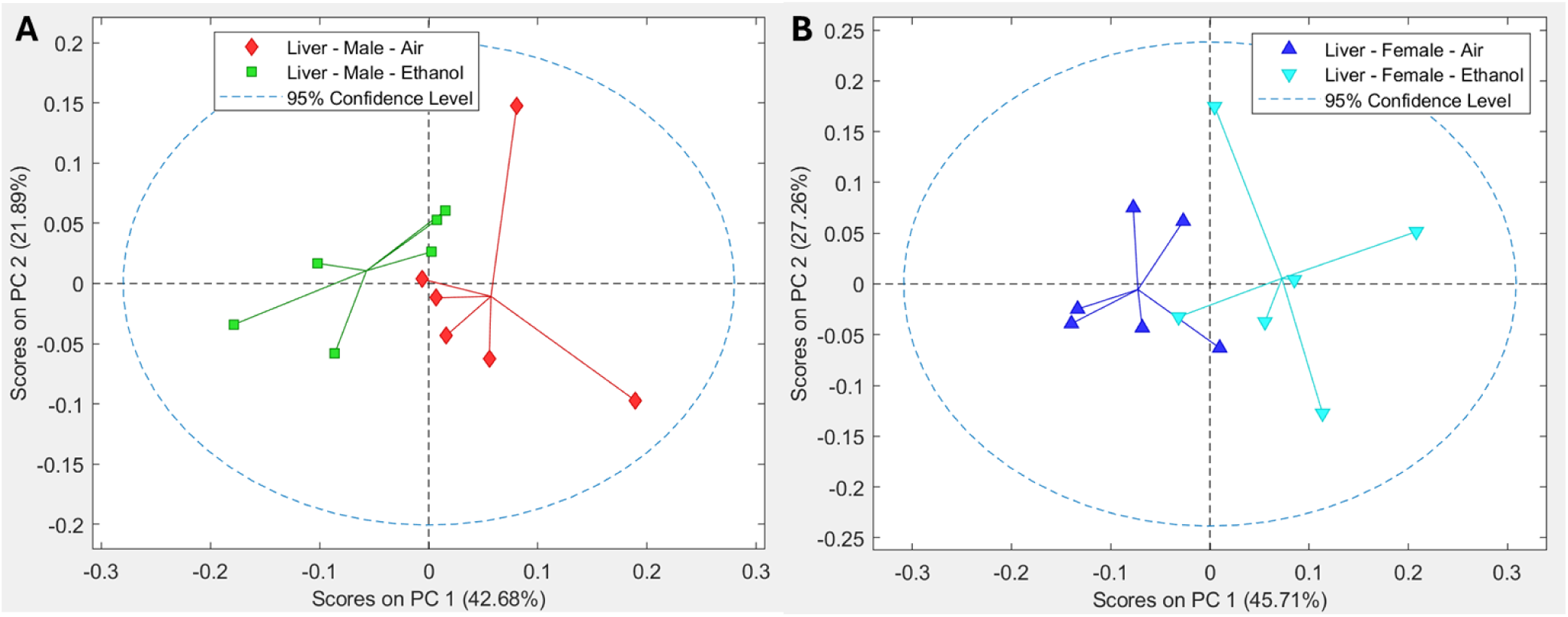
PCA studies of liver metabolites between ethanol treatment and the control. A is a score plot for the male samples and B is for the female samples. Pareto scaling was applied.

**Table 2:**
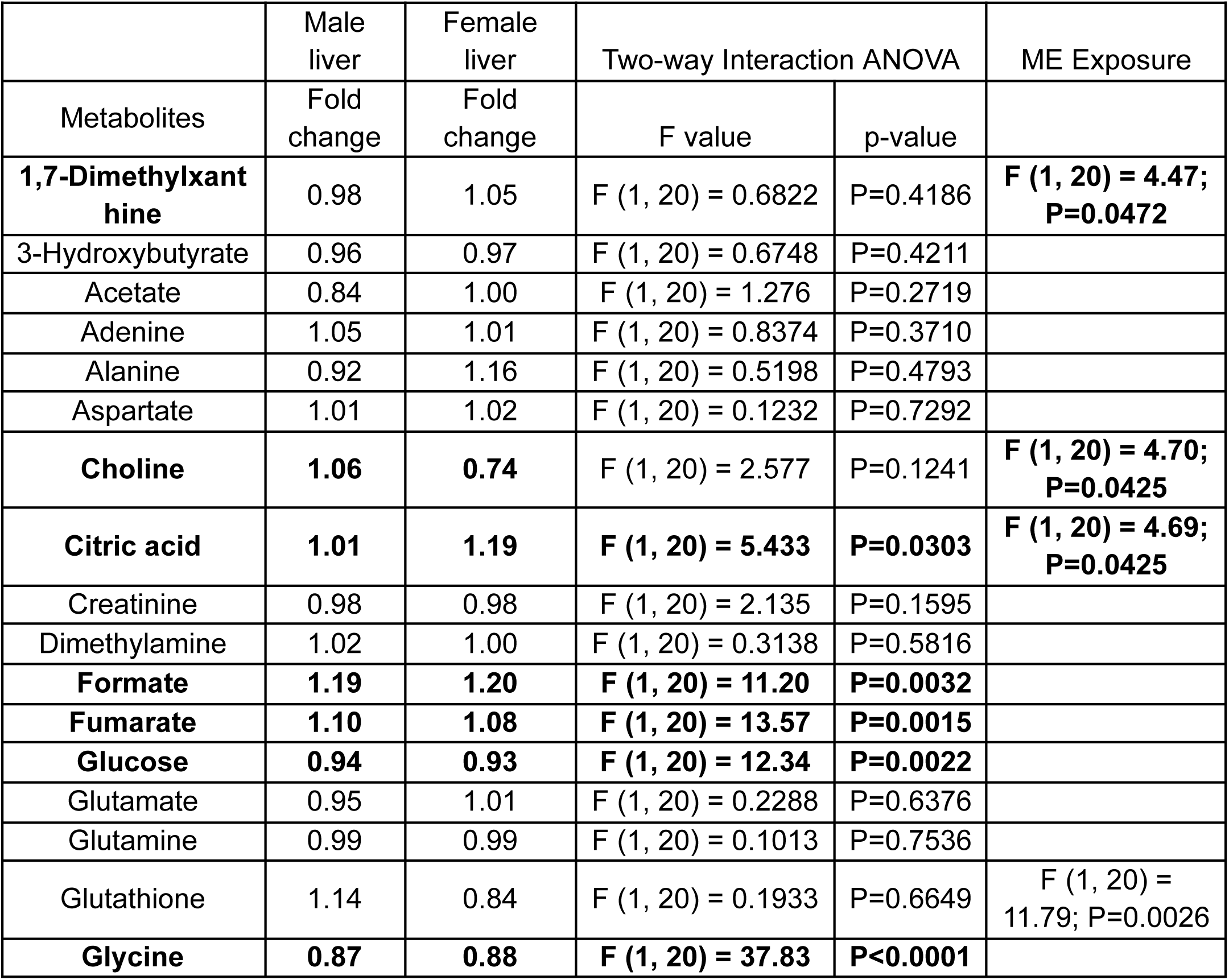
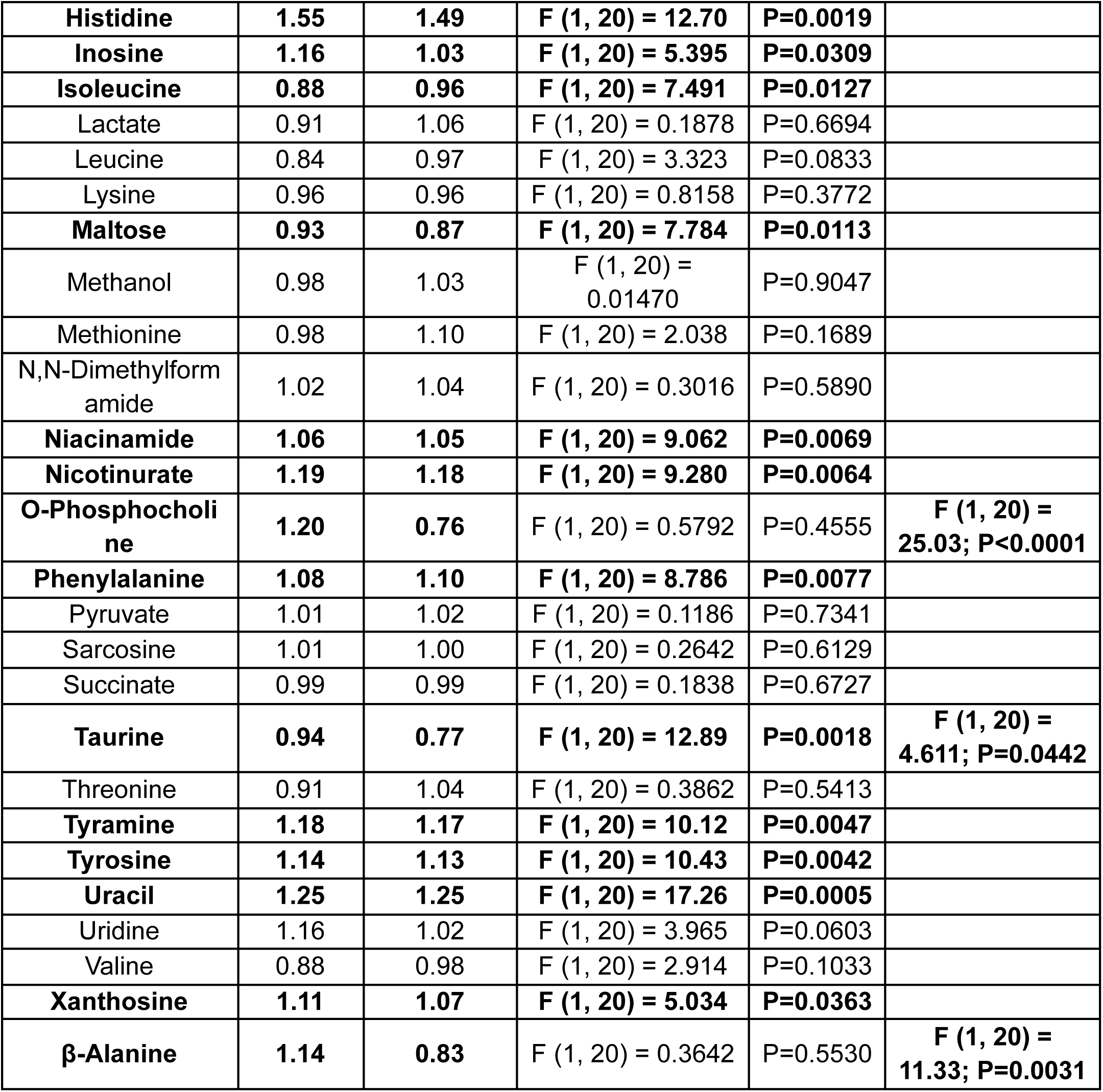
The fold change and p values of liver metabolites obtained from male and female samples. ANOVA was applied and the significant metabolite F and p-values were highlighted in bold (p<0.05).

Several metabolites are shown below in Figure 4. Individual metabolites depicted below showed statistically significant Sex by Exposure interactions (Table 2) or significant main effects of ethanol. Specifically, male CIE mice had higher niacinamide levels compared to male AIR mice (Figure 4A, p = 0.0326) and female AIR mice had higher metabolite levels compared to male AIR mice (p = 0.0272).

**Figure 4:**
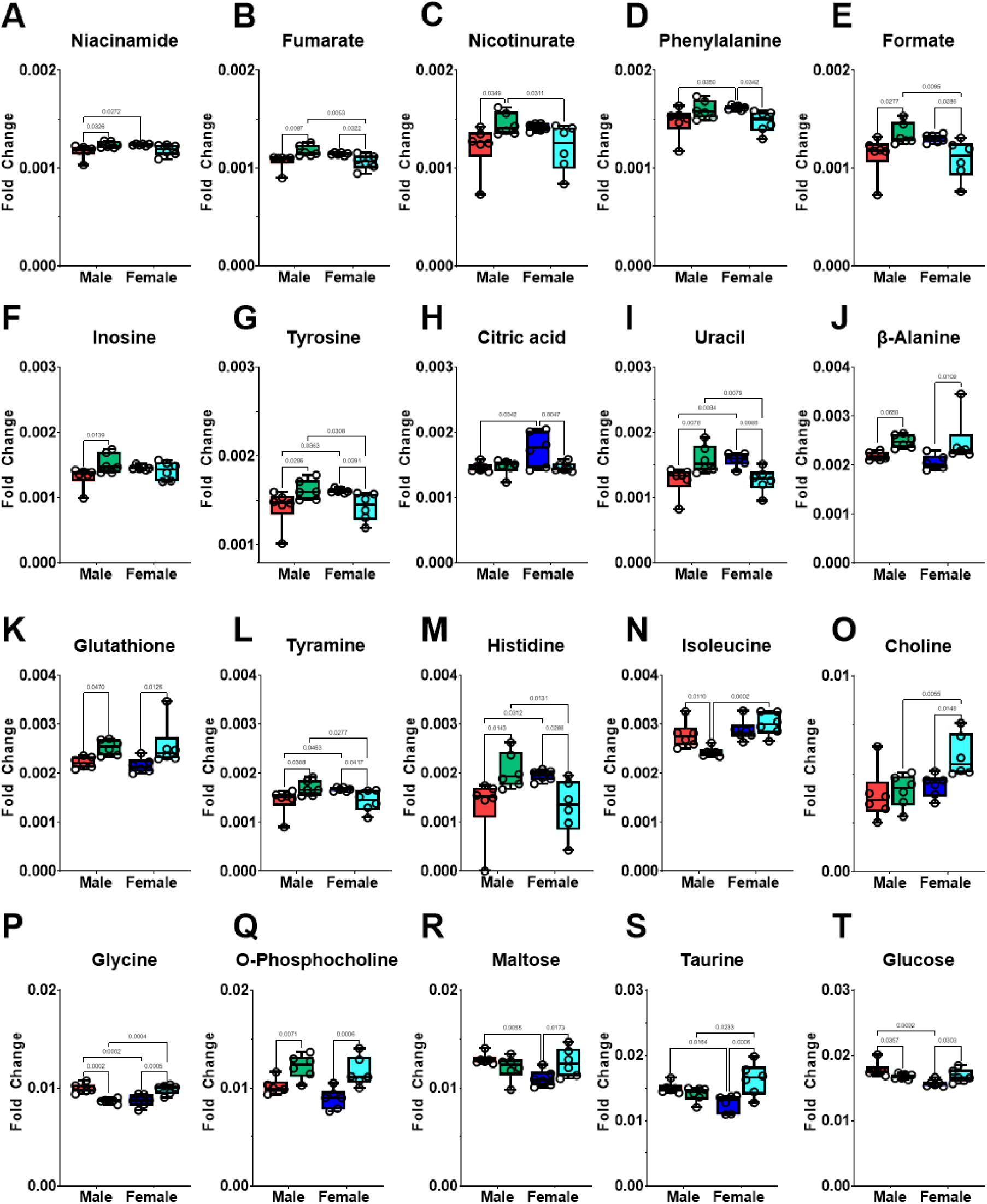
Liver metabolite comparison of individual metabolites following Chronic Intermittent Ethanol (CIE) exposure. Individual panels show the relative concentration of liver metabolites that exhibited a statistically significant Sex and Exposure interaction or a significant main effect of Exposure. Groups are color-coded as follows: Red bars represent AIR (control) males; Green bars represent CIE males; Blue bars represent AIR (control) females; and Teal bars represent CIE females. Specific p-values for all statistical comparisons are detailed in Section 3.2 and Table 2.

For fumarate (Figure 4B) and formate (Figure 4E), CIE increased metabolite levels in males (Figure 4B; p =0.0087; Figure 4E; p = 0.0277), whereas in females CIE decreased levels (p = 0.0322; p = 0.0286). Additionally, male CIE fumarate levels were higher than female CIE levels (p – 0.0053) and similarly formate (p = 0.0095). Similar patterns were observed with tyrosine (Figure 4G) with CIE increasing metabolite levels in males (p = 0.0286) and decreasing levels in females (p = 0.0391) compared to their respective controls. Additionally, male CIE mice had higher tyrosine levels compared to female CIE mice (p = 0.0308) and male AIR mice had lower tyrosine levels compared to female AIR mice (p = 0.0363). Uracil (Figure 4I) levels were increased by CIE in male mice compared to AIR (p = 0.0078) and decreased in female mice (p = 0.0085). Additionally, female AIR mice had higher uracil levels compared to male AIR mice (p = 0.0084) and female CIE mice had lower levels compared to CIE male mice (p = 0.0079). Tyramine metabolite levels (Figure 4L) were increased in CIE males compared to AIR males (p = 0.0308) and decreased in CIE females compared to their controls (p = 0.0417). CIE male tyramine levels were higher than CIE females (p = 0.0277) and lower in AIR males compared to AIR females (p = 0.0463). Histidine levels (Figure 4M) were similarly increased in male CIE mice compared to male AIR mice (p = 0.0143) and decreased in female CIE mice compared to female AIR mice (p = 0.0288). Male CIE mice also had higher histidine levels compared to female CIE mice (p = 0.0131) and male AIR levels were lower than female AIR mice (p = 0.0312). Glycine (Figure 4P_ were lower in male CIE mice compared to male AIR mice (p = 0.0002) and female CIE mice had higher metabolite levels compared to female AIR mice (p = 0.0005). In AIR mice, female glycine levels were lower than male mice (p = 0.0002) and for CIE mice, females had higher levels compared to males (p = 0.0004).

Nicotinurate levels were increased in CIE male mice compared to AIR male mice (Figure 4C; p = 0.0349) and CIE female mice (p = 0.0311). Isoleucine (Figure 4N) were lower in male CIE mice compared to male AIR mice (p = 0.0110) and female CIE mice (p = 0.0002). Choline (Figure 4O) metabolite levels were highest in female CIE mice compared to female AIR mice (p = 0.0148) and male CIE mice (p = 0.0055).

Female AIR mice had higher phenylalanine levels compared to female CIE mice (Figure 4D, p = 0.0342) and male AIR mice (p = 0.0350). Inosine metabolite levels were higher in CIE male compared to AIR male mice (p = 0.0139), whereas there were no exposure differences for female mice (Figure 4F).

For citric acid, female AIR mice had higher metabolite levels compared to female CIE mice (p = 0.0047) and male AIR mice (p = 0.0042). An inverse relationship was observed for maltose (Figure 4R) with female AIR mice having lower metabolite levels compared to female CIE mice (p = 0.0173) and male AIR mice (p = 0.0055). Similarly for taurine (Figure 4S), female AIR mice had lower levels compared to male AIR mice (p = 0.0164) and female CIE mice (p = 0.0006). Also, female CIE mice had higher taurine levels compared to male CIE mice (p 0.0233).

CIE increased β-Alanine (Figure 4J) levels in both male (p = 0.0650) and female mice (p = 0.0109), glutathione (Figure 4K; male p = 0.0470, female p = 0.0126), and O-Phosphocholine (Figure 4Q; male = 0.0071, female p = 0.0006). Finally, CIE decreased glucose (Figure 4S) levels in males compared to controls (p = 0.0357) and increased levels in females (p = 0.0303). Also, female AIR mice had lower glucose levels compared to male AIR mice (p = 0.0002).

### 3.3 Metabolomic response of serum extract after CIE

In the serum extracts, 22 metabolites were identified as shown in **Table 3**. The metabolite distributions in the PCA studies, shown in Figure 5, and the PCA score plot did not show clear separation in the male samples. However, the female samples showed partial separation. The metabolites observed were different in male and female samples. In the male samples, no significant metabolites were observed between AIR and CIE groups (p<0.05 and p<0.1). In the female samples, capric acid was significantly upregulated and methylsuccinate was significantly downregulated in CIE mice (p<0.05), while amino acids Isoleucine and tyrosine were downregulated (p<0.1) in CIE mice.

**Figure 5:**
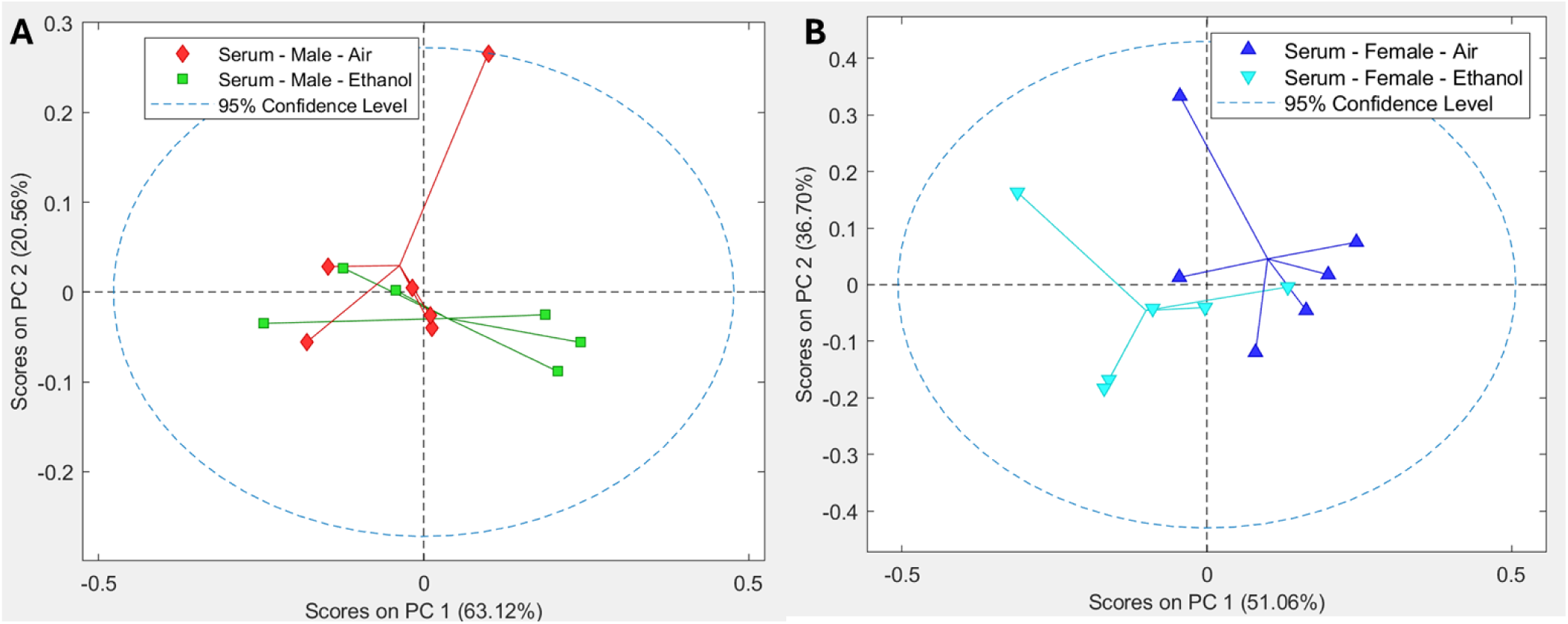
PCA score plot of the metabolites between ethanol and the control in serum samples. A is a score plot for male samples, and B is for the female samples. Pareto Scaling was applied.

**Table 3:**
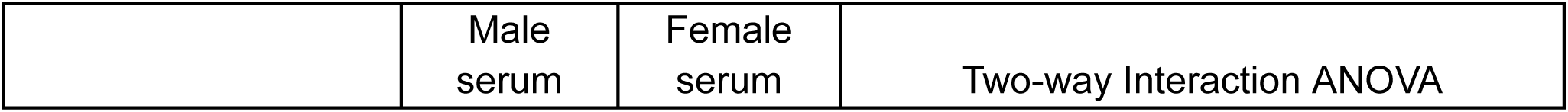
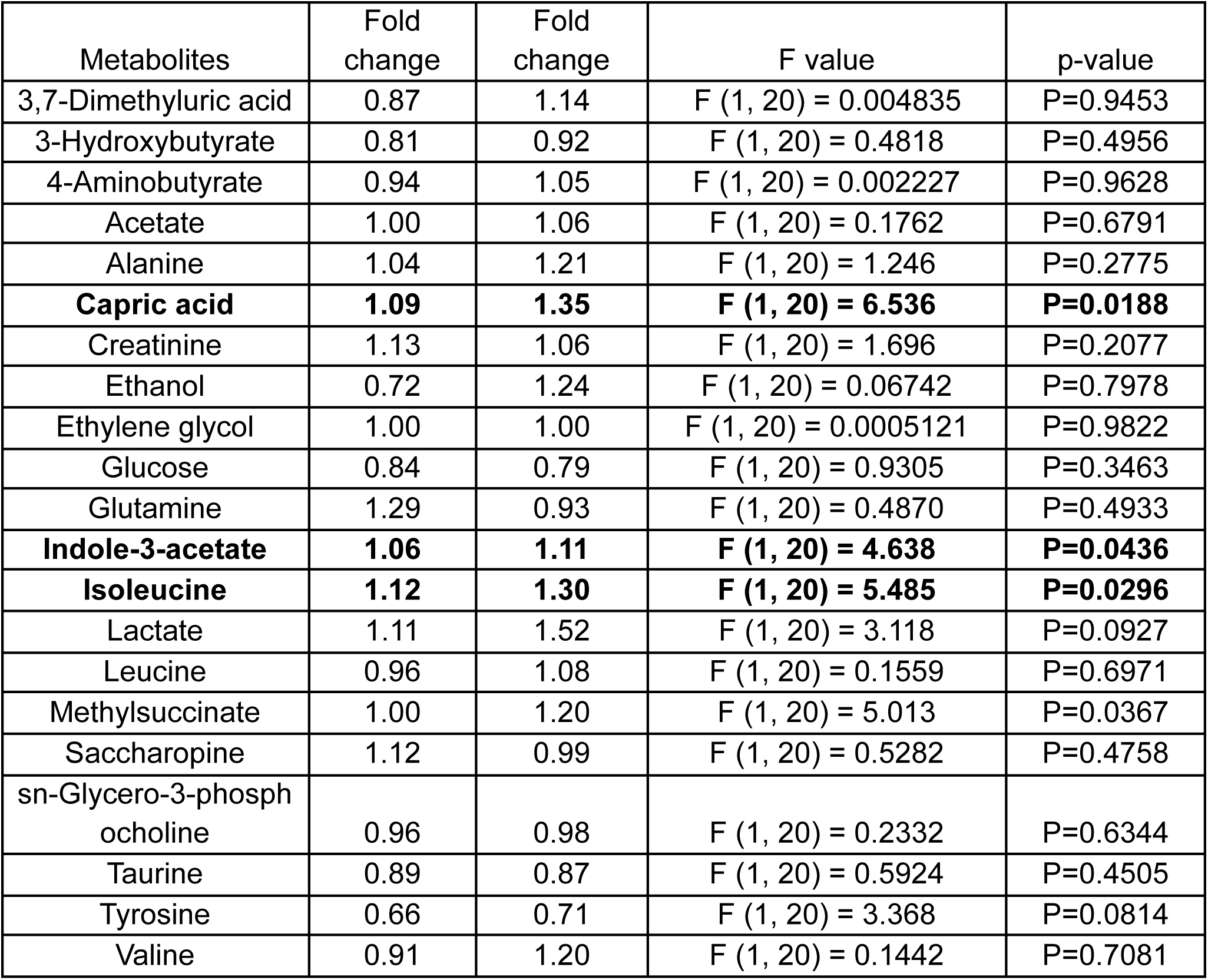
The fold change and p values of serum metabolites obtained from male and female samples. ANOVA were applied and the significant metabolite F and p-values were highlighted in bold (p<0.05).

Compared to fecal and liver samples, there were fewer sex by exposure changes in serum metabolites as shown below in Figure 6. For methylsuccinate (Figure 6A), CIE exposure decreased metabolite levels in females compared to AIE females (p = 0.0045) and there were higher metabolite levels in AIE females compared to AIE males (p < 0.0001). A similar profile was observed for Indole-3-acetate (Figure 6B) with a trend for AIR female metabolite levels higher than CIE females (p = 0.0584) and AIE males (p = 0.0083). Isoleucine (Figure 6C) showed a similar pattern with differences between AIR females and CIE females (p = 0.0234) and AIR males (p = 0.0030). Finally, capric acid (Figure 6D) levels were decreased in CIE females compared to their AIR controls (p = 0.0159) and lower than CIE males (p = 0.0002).

**Figure 6:**
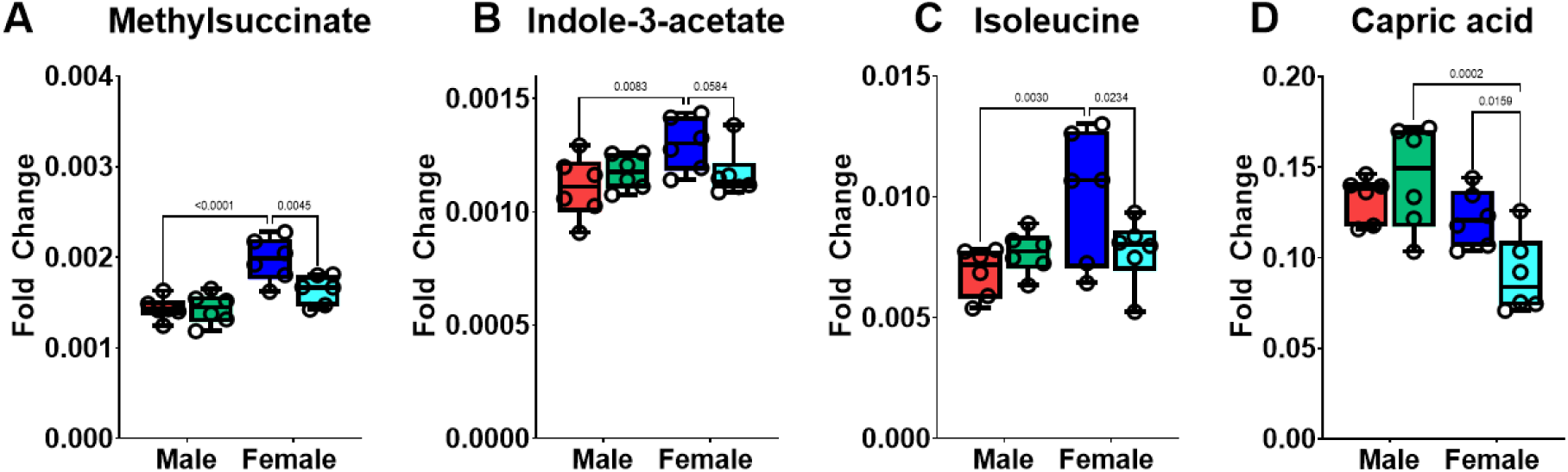
Serum metabolite comparison of individual metabolites following CIE exposure. Individual panels show the relative concentration of serum metabolites that exhibited a statistically significant Sex and Exposure interaction. Groups are color-coded as follows: Red bars represent AIR (control) males; Green bars represent CIE males; Blue bars represent AIR (control) females; and Teal bars represent CIE females. Specific p-values for all statistical comparisons are detailed in Section 3.3 and Table 3.

### 3.4 Metabolic Pathway Analysis

Pathway analyses were conducted separately for fecal, liver, and serum samples of male and female mice. Although the number of metabolites mapped for each pathway was limited, these analyses identified sex-specific pathway-level perturbations following CIE exposure.

Pathway analyses of the fecal metabolites in male mice indicated significant alterations in pyruvate metabolism, glyoxylate and dicarboxylate metabolism, and glycolysis /gluconeogenesis following CIE exposure (p<0.05; **Table 4)**. Meanwhile, in female mice, significant pathway-level changes were observed in nitrogen metabolism, arginine biosynthesis, alanine, aspartate, and glutamate metabolism, butanoate metabolism, histidine metabolism, and glutathione metabolism. Overall, the fecal pathway affected by CIE differed between male and female mice. These sex-specific fecal pathway patterns were also visualized using pathway impact plots, which show global test significance plotted against pathway impact for male and female fecal samples (Supplementary Figure S1).

**Table 4:** Selected metabolic pathways for fecal samples in male and female mice. Total compounds (Total Cmpd) refers to the total number of metabolites represented in each pathway, and Hits refers to the number of metabolites detected in the present study.

| Potential pathways | Total Cmpd | Hits | Male p-values | Female p-values |
| --- | --- | --- | --- | --- |
| Nitrogen metabolism | 6 | 2 | 2.95E-01 | **6.76E-04** |
| Arginine biosynthesis | 14 | 4 | 3.01E-01 | **7.31E-04** |
| Alanine, aspartate, and glutamate metabolism | 28 | 6 | 3.87E-01 | **2.32E-02** |
| Butanoate metabolism | 15 | 3 | 4.15E-01 | **2.57E-02** |
| Histidine metabolism | 16 | 3 | 3.18E-01 | **6.56E-04** |
| Pyruvate metabolism | 23 | 4 | **9.64E-03** | 6.82E-01 |
| Glyoxylate and dicarboxylate metabolism | 32 | 5 | **9.32E-03** | 8.03E-01 |
| beta-Alanine metabolism | 21 | 3 | 5.51E-02 | 6.56E-01 |
| Glycolysis / Gluconeogenesis | 26 | 3 | **9.64E-03** | 6.82E-01 |
| Glutathione metabolism | 28 | 3 | 5.53E-01 | **7.08E-03** |

In liver samples, pathway analysis showed substantial overlap between male and female mice. Arginine biosynthesis, glutathione metabolism, lipoic acid metabolism, porphyrin metabolism, pyrimidine metabolism, and pyruvate metabolism showed significant changes in both males and females. In contrast, nitrogen metabolism, purine metabolism, and tyrosine metabolism reached significance only in male liver samples. Alanine, aspartate and glutamate metabolism, butanoate metabolism, citrate/tricarboxylic acid cycle (TCA cycle), glycerophospholipid metabolism, and glycine, serine and threonine metabolism reached significance only in female liver samples. The liver pathway impact plots further supported the greater pathway overlap between males and females, while also showing sex-specific pathway-level patterns in liver metabolites (Figure 8).

**Figure 7:**
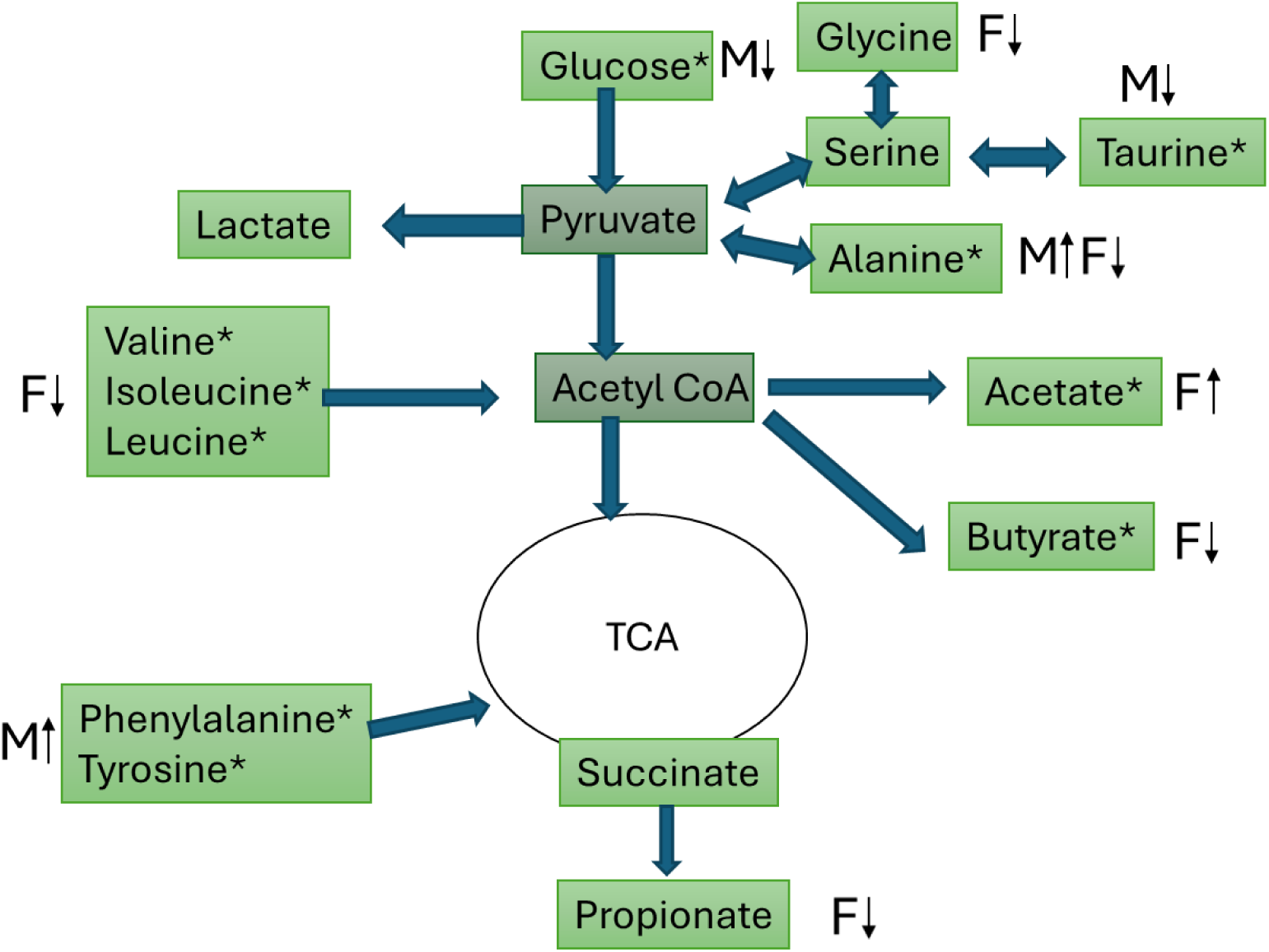
Fecal metabolites changes in general metabolic pathways for male and female mice following CIE exposure. *p<0.05, *M = is male;, and F =is female; *p<0.05 for CIE vs AIR comparisons within sex. The up arrows indicate higher are for upregulation of the specific metabolite levels in CIE mice compared to the AIR controls within the same sexgroups, and the down arrows indicate lower are for downregulation of the specific metabolite levels in CIE mice compared to the AIR controls within the same-sex groups. TCA is the tricarboxylic acid cycle.

**Figure 8:**
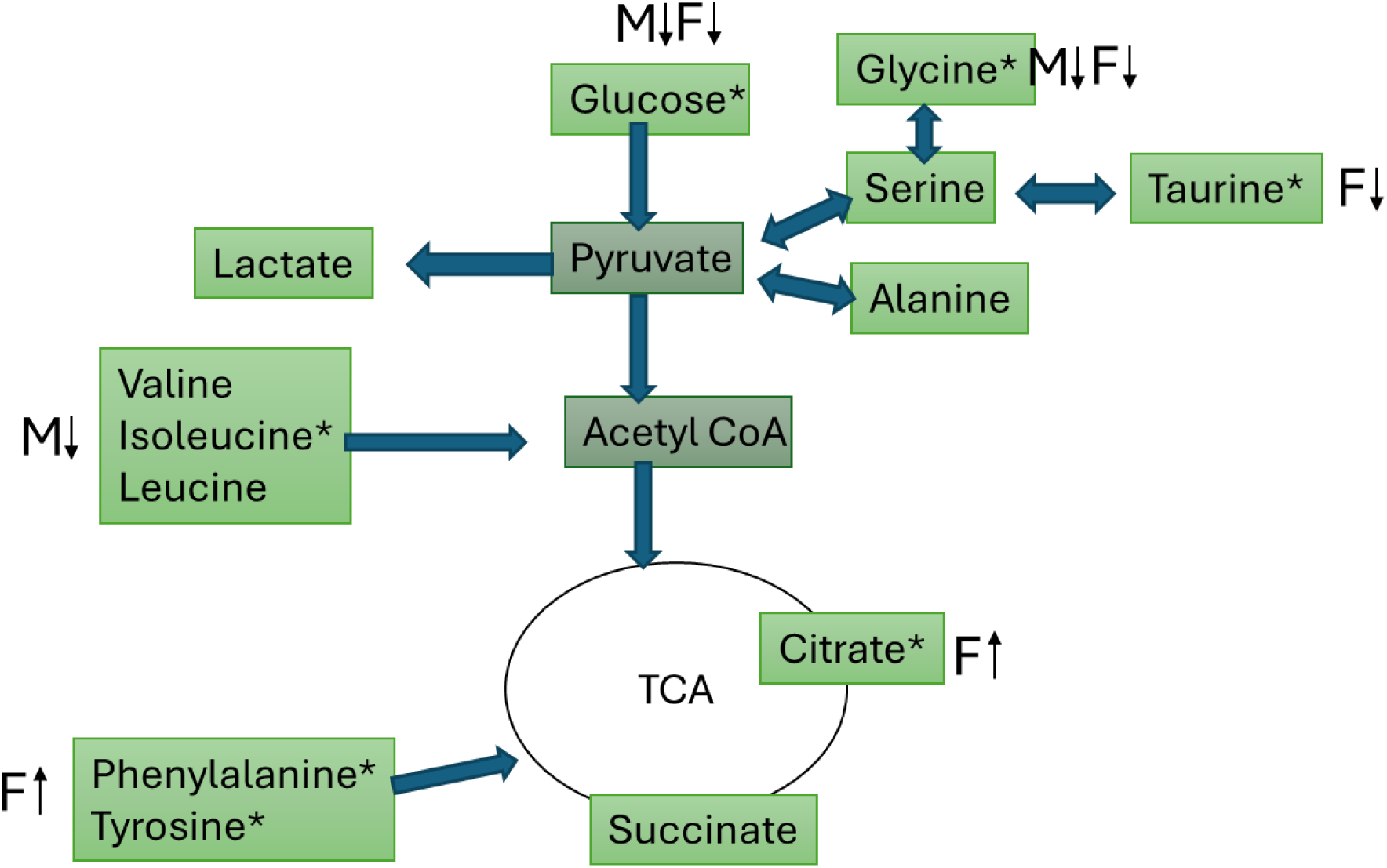
Liver metabolite changes mapped onto major metabolic pathways in male and female mice following CIE. M = male; F = female; *p<0.05 for CIE vs AIR comparisons within sex, Up arrows indicate higher metabolite levels in CIE mice relative to AIR controls within the same sex, and down arrows indicate lower metabolite levels in CIE mice relative to AIR controls within the same sex. TCA = tricarboxylic acid cycle.

In serum samples, no pathways reached statistical significance in either male or female mice (**Table 6**), consistent with the limited number of significant metabolites observed.

**Table 5:** Selected metabolic pathways identified from liver samples in male and female mice. Total compounds (Total Cmpd) refers to the total number of metabolites represented in each pathway, and Hits refers to the number of metabolites detected in the present study.

|  | Total Cmpd | Hits | Male p-values | Female p-values |
| --- | --- | --- | --- | --- |
| Alanine, aspartate and glutamate metabolism | 28 | 6 | 6.60E-02 | **1.29E-02** |
| Arginine biosynthesis | 14 | 4 | **4.68E-02** | **4.76E-02** |
| Butanoate metabolism | 15 | 2 | 1.32E-01 | **1.19E-02** |
| Citrate cycle (TCA cycle) | 20 | 3 | 7.19E-02 | **1.35E-02** |
| Glutathione metabolism | 28 | 3 | **6.20E-04** | **3.74E-03** |
| Glycerophospholipid metabolism | 36 | 2 | 7.11E-01 | **2.12E-02** |
| Glycine, serine and threonine metabolism | 34 | 5 | 3.19E-01 | **1.78E-02** |
| Lipoic acid metabolism | 28 | 2 | **2.84E-02** | **3.92E-02** |
| Nitrogen metabolism | 6 | 2 | **3.39E-03** | 5.65E-02 |

|  | Total<br>Cmpd | Hits | Male<br>p-values | Female<br>p-values |
| --- | --- | --- | --- | --- |
| Alanine, aspartate and glutamate metabolism | 28 | 6 | 6.60E-02 | **1.29E-02** |
| Arginine biosynthesis | 14 | 4 | **4.68E-02** | **4.76E-02** |
| Butanoate metabolism | 15 | 2 | 1.32E-01 | **1.19E-02** |
| Citrate cycle (TCA cycle) | 20 | 3 | 7.19E-02 | **1.35E-02** |
| Glutathione metabolism | 28 | 3 | **6.20E-04** | **3.74E-03** |
| Glycerophospholipid metabolism | 36 | 2 | 7.11E-01 | **2.12E-02** |
| Glycine, serine and threonine metabolism | 34 | 5 | 3.19E-01 | **1.78E-02** |
| Lipoic acid metabolism | 28 | 2 | **2.84E-02** | **3.92E-02** |
| Phenylalanine metabolism | 10 | 2 | 5.30E-02 | 2.39E-01 |
| Phenylalanine, tyrosine, and tryptophan biosynthesis | 4 | 2 | 5.30E-02 | 2.39E-01 |
| Porphyrin metabolism | 31 | 2 | **2.86E-02** | **3.96E-02** |
| Purine metabolism | 71 | 4 | **2.31E-04** | 5.05E-02 |
| Pyrimidine metabolism | 39 | 3 | **1.68E-03** | **4.42E-02** |
| Pyruvate metabolism | 23 | 4 | **2.58E-02** | **4.40E-02** |
| Tyrosine metabolism | 42 | 4 | **4.96E-02** | 5.91E-02 |

**Table 6:** The selected metabolomics pathways for serum samples. Fold change metric was calculated using the average of each metabolite concentration in the treated group divided by the average of each metabolite concentration in the control group

|  | Male liver |  | Female liver |  |
| --- | --- | --- | --- | --- |
| Metabolites | p-value | Fold change | p-value | Fold change |
| O-Phosphocholine | 6.64E-03 | 1.2 | **4.25E-03** | 0.76 |
| Uracil | 2.96E-02 | 1.25 | **4.53E-03** | 1.25 |
| Glycine | 1.66E-04 | 0.87 | **5.51E-03** | 0.88 |
| Taurine | 1.32E-01 | 0.94 | **9.60E-03** | 0.77 |
| Choline | 7.28E-01 | 1.06 | **1.23E-02** | 0.74 |
| Phenylalanine | 1.42E-01 | 1.08 | **1.24E-02** | 1.1 |
| Tyrosine | 7.20E-02 | 1.14 | **1.91E-02** | 1.13 |
| Histidine | 3.85E-02 | 1.55 | **2.02E-02** | 1.49 |
| Fumarate | 2.76E-02 | 1.1 | **2.24E-02** | 1.08 |
| Tyramine | 7.13E-02 | 1.18 | **2.45E-02** | 1.17 |
| Formate | 5.01E-02 | 1.19 | **2.92E-02** | 1.2 |
| Citric acid | 8.56E-01 | 1.01 | 3.07E-02 | 1.19 |
| Glucose | 5.42E-02 | 0.94 | 3.64E-02 | 0.93 |
| Maltose | 1.63E-01 | 0.93 | 3.90E-02 | 0.87 |
| Methionine | 7.22E-01 | 0.98 | 4.17E-02 | 1.1 |
| Niacinamide | 6.67E-02 | 1.06 | 4.67E-02 | 1.05 |
| 1,7-Dimethylxanthine | 4.11E-01 | 0.98 | 5.03E-02 | 1.05 |
| Nicotinurate | 5.79E-02 | 1.19 | 5.48E-02 | 1.18 |
| Glutathione | 2.76E-03 | 1.14 | 6.28E-02 | 0.84 |
| β-Alanine | 1.13E-03 | 1.14 | 6.38E-02 | 0.83 |
| Creatinine | 4.39E-01 | 0.98 | 9.29E-02 | 0.98 |
| Alanine | 1.74E-01 | 0.92 | 1.68E-01 | 1.16 |
| Xanthosine | 1.11E-01 | 1.11 | 1.91E-01 | 1.07 |
| Isoleucine | 1.37E-02 | 0.88 | 3.38E-01 | 0.96 |
| Inosine | 4.71E-02 | 1.16 | 4.56E-01 | 1.03 |

## 4 Discussion

The main finding from this study is that CIE did not produce one uniform metabolic profile across sample types or between males and females. Using adult male and female C57BL/6J mice, we assessed ethanol-associated metabolic changes three days after the final AIR or CIE exposure cycle across fecal, liver, and serum samples. Our results demonstrated that metabolic pathway perturbations were most pronounced in fecal and liver samples, while serum showed fewer metabolite and pathway-level changes. Moreover, the metabolic pathways affected by CIE differed between males and females, indicating that the metabolic response to CIE was both sex-specific and sample-type specific. These findings support the translational value of metabolomics for identifying tissue– and sex-specific metabolic signatures that may help explain individual vulnerability to alcohol-related health outcomes (Nicholas et al., 2008; Mamas et al., 2011; Clish, 2015; Kezer et al., 2021; Bizzaro et al., 2023).

### 4.1 Fecal metabolites

The PCA score plots showed partial separation between AIE and CIE groups within male and female fecal samples (Figure 1). This visual separation was based on the relative clustering of AIR and CIE samples in the PCA score plots, with group centers appearing more separated in females than males in the PC1 direction. PCA interpretation was guided by the approach described in Mayonu and colleagues with good separation (p < 0.05) (Mayonu et al., 2026).

Pathway analysis showed that fecal metabolic pathways differed between males and females following CIE exposure (Xia and Wishart, 2010). Because the number of detected metabolites contributing to each pathway was limited, these results should be interpreted as candidate pathway-level differences rather than definitive evidence of broad pathway disruption. However, these sex-specific pathway patterns suggest that CIE altered fecal metabolic profiles differently in males and females. The metabolite-level findings are discussed below, with emphasis on female-specific short-chain fatty acids(SCFAs) and branched-chain amino acids (BCAAs), as well as male-specific changes in gut-associated metabolites (Morrison and Preston, 2016; Gojda and Cahova, 2021).

Female fecal metabolites showed SCFAs changes following CIE, including lower butyrate, increased acetate, and a trend toward lower propionate in females (Figure 2, **Table 1**). Because these SCFAs are largely produced by gut bacteria during fermentation (Imdad et al., 2022) and reflect gut-derived metabolite output (Morrison and Preston, 2016; Deleu et al., 2021), this pattern suggests that CIE altered the fecal metabolic profile in females. Increased acetate combined with decreased butyrate is important because acetate can contribute to butyrate formation by fecal bacteria, so changes in the opposite direction may reflect a shift in fermentation balance (Duncan et al., 2004; LaBouyer et al., 2022). Butyrate supports colonic health and intestinal function (Deleu et al., 2021; Stoeva et al., 2021; Wijdeveld et al., 2023). The propionate finding is also important because previous mouse work showed that propionate reduced alcohol-induced liver injury through gut-liver axis mechanisms (Xu et al., 2022). In the present study, propionate trended lower in CIE females, suggesting that CIE may reduce a SCFA that has been linked to protection in alcohol-related liver injury models. We did not measure liver injury, microbiome composition, or intestinal permeability in the present work, so this should be treated as a candidate mechanism for following up rather than direct evidence of liver damage (Hartmann et al., 2015; Szabo, 2015).

Branched-chain amino acids (BCAAs), including leucine, isoleucine, and valine were all reduced in female fecal samples following CIE exposure. This coordinated BCAA decrease was strongest in the fecal samples and was not mirrored as a clear female BCAA pattern in liver or serum. Chronic ethanol exposure alters gastrointestinal metabolites, fecal metabolite profiles, and gut microbial composition in alcohol-related models, including those of alcohol-associated fatty liver disease (Xie et al., 2013; Shi et al., 2015; Piacentino et al., 2021). BCAAs can reflect diet, host metabolism, and microbial metabolism, so fecal metabolomics alone cannot identify the exact source of the decrease (Gojda and Cahova, 2021). Here, the decrease in leucine, isoleucine, and valine suggests that CIE altered fecal amino-acid related metabolites in females, adding to the fecal SCFA pattern described above.

In males, CIE reduced fecal taurine and glucose compared to AIR controls. Taurine has been linked to alcohol-related liver and oxidative outcomes in humans. In chronic alcoholic patients, administration of taurine reduced AST, ALT, bilirubin, and TBARS, consistent with lower liver-cell injury markers and reduced oxidative lipid damage (Hsieh et al., 2014). Taurine could reduce fatty degeneration and lipid accumulation in the liver caused by alcohol consumption (Wu et al., 2015). In rat models of chronic ethanol exposure, taurine reduced ethanol-induced hepatic steatosis, lipid accumulation, lipid peroxidation, and hepatic inflammation (Kerai et al., 1998; Lin et al., 2015; Wu et al., 2015). In the present work, the combined decrease in fecal taurine and glucose identifies a male-specific ethanol-associated fecal metabolite pattern. This pattern is consistent with prior work showing that chronic ethanol exposure alters gastrointestinal metabolites and gut-liver signaling pathways (Xie et al., 2013; Szabo, 2015; Albillos et al., 2020).

In addition, in the present work CIE increased fecal phenylalanine and tyrosine in males, which may reflect altered aromatic amino acid metabolism (Liu et al., 2020; Zhang et al., 2022; Mrdjen et al., 2023). Altered aromatic amino acid profiles have been reported in alcohol-liver disease, and chronic alcohol-associated metabolome studies have identified disruptions in amino acid-related pathways (Liu et al., 2020). Human studies of alcohol-related liver cirrhosis have also identified altered fecal and plasma amino acid metabolites, including phenylalanine and tyrosine (Xu et al., 2023). Unlike females, males showed fewer significant SCFA shifts, suggesting a different pattern of gut-associated fecal metabolic response rather than the same SCFA/BCAA-heavy pattern observed in females (Morrison and Preston, 2016; Gojda and Cahova, 2021). Together, the male fecal profile was marked by lower taurine and glucose and higher phenylalanine and tyrosine, supporting a sex-specific fecal metabolite response to CIE.

### 4.2 Liver metabolites

While the pathway analysis revealed more overlap between males and females in liver samples than was observed in fecal samples, the individual metabolite shifts were still informative. Glucose is central to hepatic energy metabolism, and the liver regulates systemic glucose homeostasis through pathways that include glycogen storage, glycolysis, and gluconeogenesis (Sharabi et al., 2015; Han et al., 2016; Nakrani et al., 2020). In this study, glucose decreased in CIE-exposed females compared to AIR females and trended lower in CIE-exposed males compared with AIR males. The liver plays a major role in glucose homeostasis and systemic energy balance (Sharabi et al., 2015; Han et al., 2016). Glycine was reported as a therapeutic immunonutrient in alcohol-related liver disease, with proposed anti-inflammatory effects that may reduce alcohol-related hepatic injury (Yamashina et al., 2005; Aguayo-Cerón et al., 2023). Together, decreased liver glucose and glycine suggest that CIE altered hepatic metabolites linked to energy balance and alcohol-related liver vulnerability in both males and females (Yamashina et al., 2005; Sharabi et al., 2015; Aguayo-Cerón et al., 2023). Because liver injury endpoints were not measured here, these findings should be interpreted as metabolomic evidence of an altered hepatic metabolic state rather than direct evidence of liver disease (Nicholas et al., 2008; Mamas et al., 2011).

In female liver samples, CIE decreased taurine and increased citric acid, phenylalanine, and tyrosine compared with AIR controls. Taurine is reduced in several liver disease states and contributes to bile acid conjugation, which supports lipid digestion and absorption (Miyazaki and Matsuzaki, 2014; Wang et al., 2024). Interestingly, reduced taurine and increased levels of phenylalanine and tyrosine were observed in female liver samples and in male fecal samples. This overlap suggests that CIE affected similar metabolite classes across different biological compartments, but not in the same sex or tissue. Male liver samples did not show the same phenylalanine and tyrosine pattern observed in female liver samples. Instead, the male aromatic amino acid response was more apparent in fecal samples, suggesting a tissue-specific CIE effect that may involve gut-associated aromatic amino acid metabolism (Liu et al., 2020; Mrdjen et al., 2023). Overall, the female liver profile suggests that CIE altered hepatic metabolites linked to taurine and bile acid biology, energy metabolism, and aromatic amino acid metabolism (Miyazaki and Matsuzaki, 2014; Han et al., 2016; Mrdjen et al., 2023). Because liver injury endpoints were not measured here, these findings should be interpreted as metabolomic evidence of a female-biased hepatic response to CIE rather than direct evidence of liver injury (Nicholas et al., 2008; Mamas et al., 2011).

## 5 Conclusions

This study examined the sex– and tissue-specific metabolomic alterations in adult C57BL/6J mice three days after CIE exposure across fecal, liver, and serum samples. Serum metabolite changes were limited, which may reflect the three-day recovery period following CIE exposure. The strongest ethanol-associated fecal changes were observed in females. Several key short-chain fatty acids (SCFAs) including acetate, butyrate, and propionate, were altered suggesting that CIE influences microbial metabolism and gut-associated energy substrates. In male fecal samples, reduced taurine and glucose levels were consistent with a different ethanol-associated gut-metabolite profile.. In liver tissues, decreased glucose and glycine in both males and females suggested ethanol-associated shifts in hepatic energy metabolism.

Notably, taurine, phenylalanine, and tyrosine showed changes in male fecal samples and in female liver samples. Together, these findings suggest that CIE did not produce one uniform metabolic response. Instead, the metabolite patterns differed by sex and sample type, with females showing strong fecal SCFA/BCAA changes and a distinct liver metabolite profile, while males showed a fecal profile marked by taurine, glucose, phenylalanine, and tyrosine. Because liver injury endpoints such as ALT, AST, or histology were not assessed, these results should be interpreted as metabolomic signatures of potential vulnerability rather than confirmed liver damage.

Collectively, these results provide evidence for sex-specific metabolic effects of CIE and support the identification of candidate metabolite signatures. Future studies should integrate metabolomics with direct microbiome profiling, such as 16S rRNA sequencing and/or metagenomics. Additional work should extend analyses to brain and heart tissue following CIE exposure. Ultimately, these findings may inform future mechanistic studies and therapeutic approaches for alcohol-related disorders.

## 7 Declarations

The authors declare that the research was conducted in the absence of any commercial or financial relationships that could be construed as a potential conflict of interest. All authors have read the version and approved it to be published.

## 8 Data availability

The raw data supporting the conclusions of this article will be made available by the authors without undue reservation.

## 7 Supporting materials

**Figure 9:**
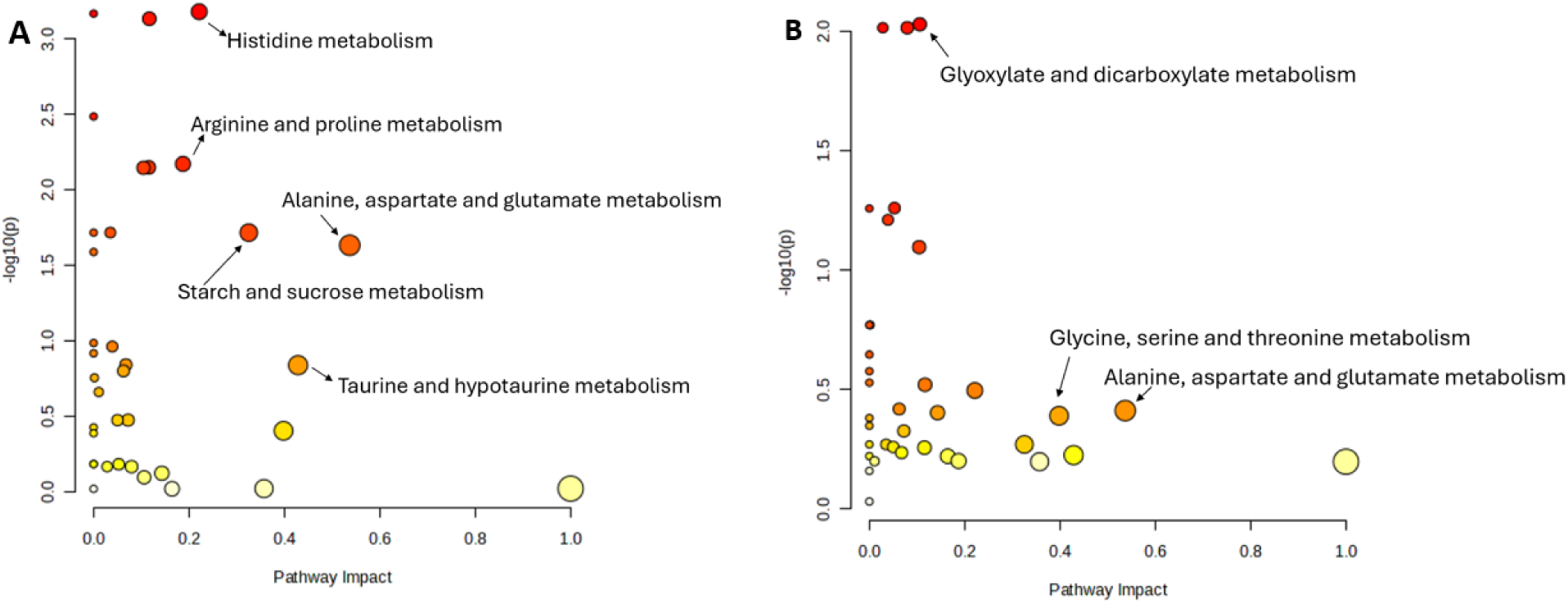
The fecal metabolic pathway analysis using the plot of the global t-test vs. the pathway impact. A, male, and B, female.

**Figure 10:**
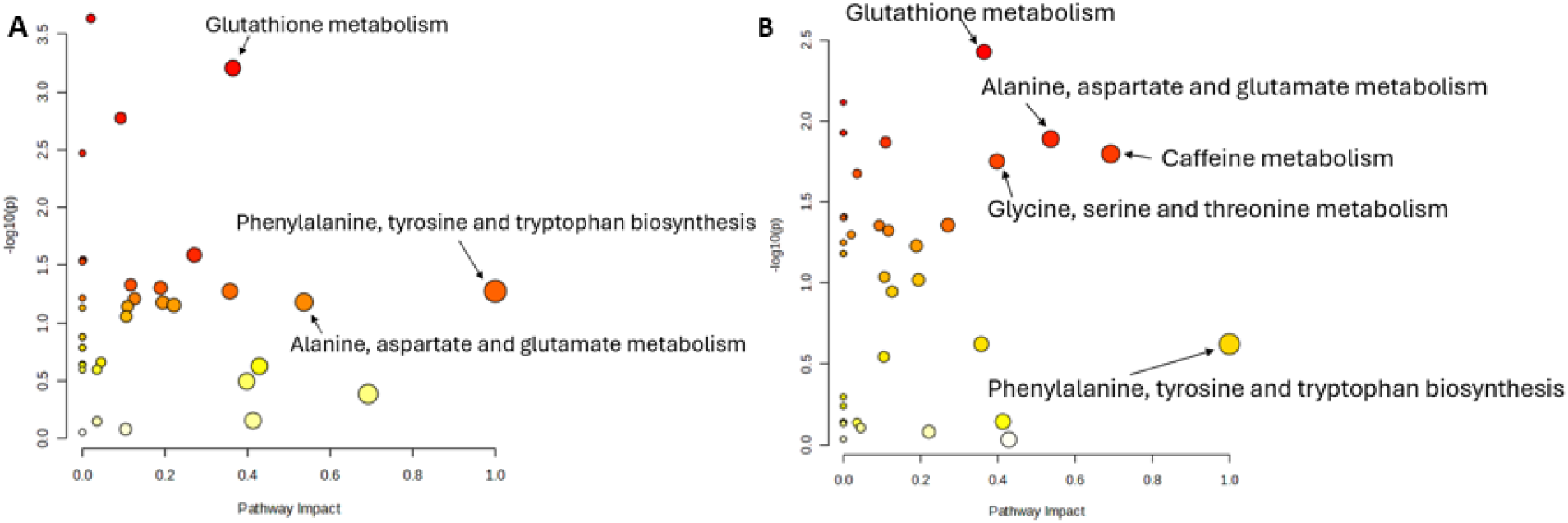
The liver metabolic pathway analysis using the plot of the global t-test vs. the pathway impact. A, male, and B, female.

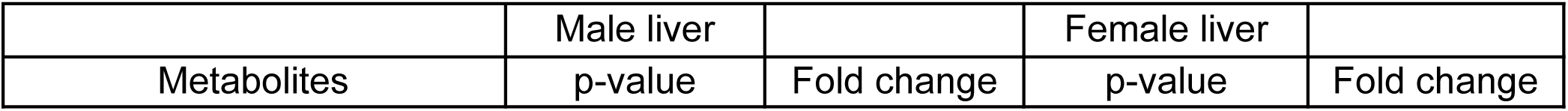
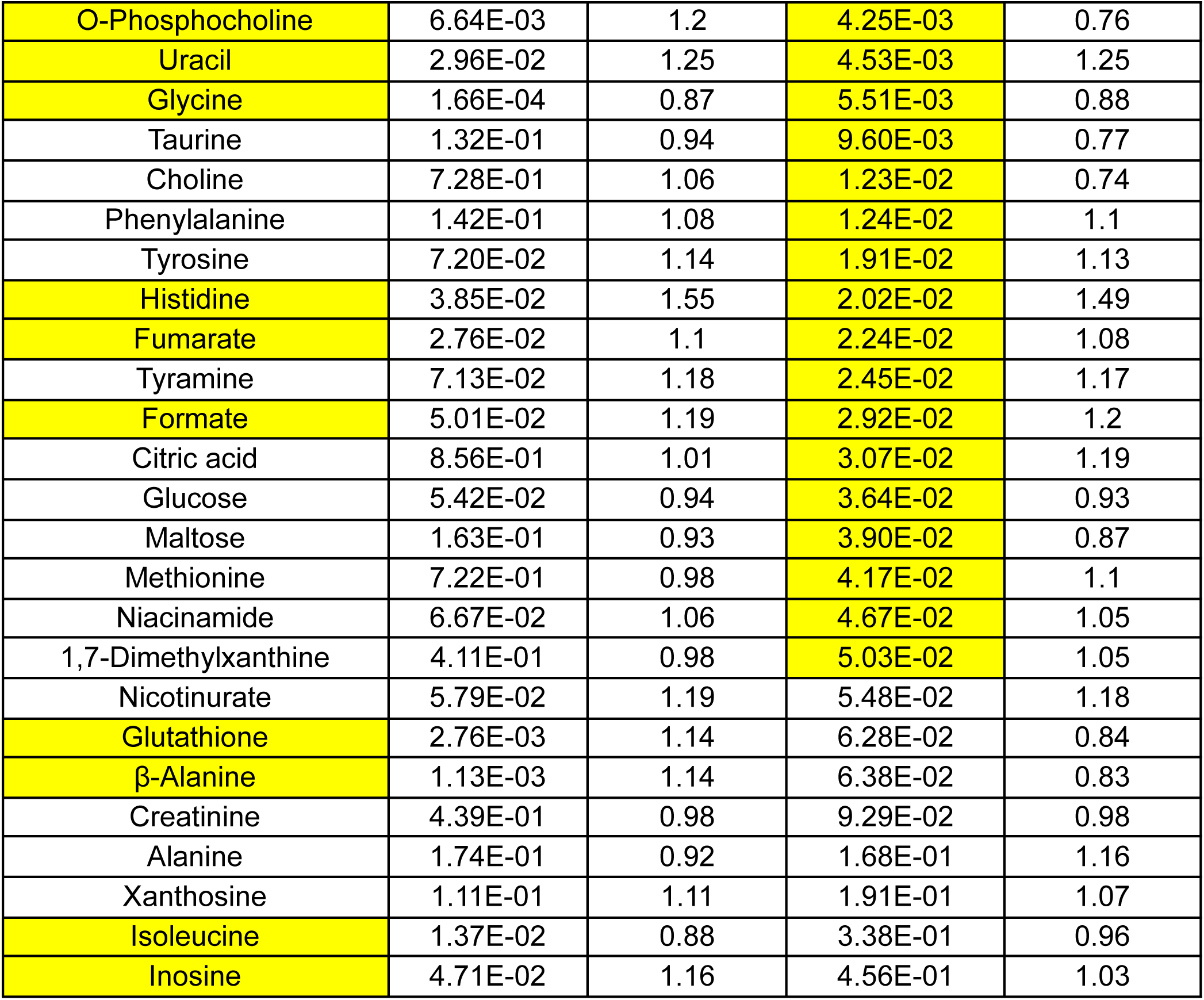
Table 6:

